# An association between sexes of successive siblings in the data from Demographic and Health Survey program

**DOI:** 10.1101/031344

**Authors:** Mikhail Monakhov

**Keywords:** sex ratio, sex determination, Lexian variation, Markov dependency, sex composition

## Abstract

The prediction of future child’s sex is a question of keen public interest. The probability of having a child of either sex is close to 50%, although multiple factors may slightly change this value. Some demographic studies suggested that sex determination can be influenced by previous pregnancies, although this hypothesis was not commonly accepted. This paper explores the correlations between siblings’ sexes using data from the Demographic and Health Survey program. In the sample of about 2,214,601 women (7,985,855 children), the frequencies of sibships with multiple siblings of the same sex were significantly higher than can be expected by chance. A formal modelling demonstrated that sexes of the children were dependent on three kinds of sex ratio variation: a variation between families (Lexian), a variation within a family (Poisson) and a variation contingent upon the sex of preceding sibling (Markovian). There was a positive correlation between the sexes of successive siblings (coefficient = 0.067, p < 0.001), i.e. a child was more likely to be of the same sex as its preceding sibling. This correlation could be caused by secondary sex ratio adjustment *in utero* since the effect was decreasing with the length of birth-to-birth interval, and the birth-to-birth interval was longer for siblings with unlike sex.

## Introduction

A family with multiple children of same sex, say, a family with ten boys and no girls, always draws attention, and speculations are made about possible reasons of such an incident. These may include a lack of either parent’s ability to conceive girls, an influence of witchcraft, astrology etc.

Surely, large unisex families may emerge simply by chance, when sexes of siblings are completely independent from each other. However, some studies indicate that the sex of a child may depend statistically on sexes of other siblings (see references below). There are at least two possible sources of such dependence, that may increase the frequency of unisex families. First, some parents indeed may have a predisposition to have children of particular sex. This bias can be caused by chemical compounds, infectious diseases, psychological distress (James 2000) and other conditions specific for given parents. As a result, each couple can be characterized by a probability of male birth *p* that may differ from the frequency of boys in the population^1^. This variation of *p* among couples is dubbed “Lexian”.

Second, the sex of a child could be affected somehow by the sex of immediately preceding sibling, or sexes of several preceding siblings. This effect could be positive (the birth of a child with either sex increases chances that next child will be of same sex) or negative. The variation of *p* within a sibship, dependent on previous birth, is called “Markovian”. The unisex families would be in excess in a population with positive Markovian correlation.

Another source of variation in sex ratio is called “Poisson”. It refers to random or systematic variation of *p* within a family, for example a decrease of *p* with mother’s age. Markovian, Lexian and Poisson variations can be present simultaneously in the same sample. Additionally, statistical dependence between siblings’ sexes can be caused by parents’ sex preferences, e.g. boy preference documented in many Asian cultures, or a preference for mixed-sex sibship.

Since at least 1889^2^ the problem was studied using demographic data. Geissler ((Geissler 1889), reviewed in (Gini 1951)) analyzed about 5 million births in 19^th^ century Germany and did not detect significant deviation from expected sibship frequencies. Nevertheless, he noted that in same-sex sibships probability of birth of one more child of same sex was higher than in other families (a positive correlation). For mixed sex sibships the correlation was negative (for instance, probability of female birth increased after many male births).

Gini (Gini 1951) re-analyzed Geissler’s data, together with additional datasets from Germany, Italy and Netherlands, and concluded that some couples had a predisposition to have children of particular sex, although this predisposition may reverse with time. For example, a family may start procreation with a tendency to produce one sex, and end with a tendency to produce the opposite sex. He noticed a positive association between sexes of successive siblings in a data from Italy, and interpreted it as an evidence for such predispositions.

Turpin and Schutzenberger (Turpin and Schutzenberger 1949) (reviewed in (Gini 1951)) analyzed a sample from France (14,230 families) and observed positive correlation between sexes of successive siblings. Unlike Gini, they hypothesized that the correlation could be explained by an influence of one birth on the following birth. This idea was supported by the fact that the interval between births of same sex siblings was on average shorter than for opposite sex siblings (however, Gini interpreted this result in favor of his “reversal of predisposition” explanation).

(Bernstein 1952) analyzed a sample from USA (7,616 families). She reported positive correlation between sexes of first two children in family, and an excess of unisexual three-child families.

Malinvaud (Malinvaud 1955) (reviewed in (James 1975)) investigated nearly 4 million births in France. The probability of male birth correlated positively with number of preceding boys and negatively – with number of preceding girls.

Renkonen *et al* (Renkonen 1956, Renkonen, Makela et al. 1961) collected data from 31215 families in Finland. Their dataset was analyzed by several researchers, who reached contradictory conclusions. Original authors confirmed associations reported earlier by Geissler (Renkonen, Makela et al. 1962). First, probability of male birth was higher when all previous siblings were boys, and lower when they were girls (positive correlation). Second, for mixed sibships the probability of male birth was decreasing when number of preceding boys increased (negative correlation). The negative correlation was explained by possible immunization of mother’s organism against male fetal antigens. Edwards (Edwards 1961, Edwards 1962, Edwards 1966) re-analyzed the data using different statistical methods and demonstrated that correlation between sexes of successive siblings was positive (presumably due to some kind of *in utero* adjustment of sex ratio), although it was negative when only two last siblings in a family were concerned (probably due to birth control). Beilharz (Beilharz 1963) analyzed same data and dismissed previous conjectures about biological factors. Instead, he concluded that there is no influence of sex of previous sibling, and apparent positive correlation is caused by heritable tendency of some parents to have same-sex sibships.

No correlation was found between successive siblings’ sexes in 116,458 sibships from Utah, USA (Greenberg and White 1967). Also, interval between births in this sample did not depend on whether the siblings were of same or opposite sex.

In the data from 1970 USA census, with more than 230,000 families, sex ratio increased with number of preceding boys, and decreased with number of preceding girls. The authors suggested that their data provide evidence for both Markovian and Lexian variation (Ben-Porath and Welch 1976).

In another USA sample (649,366 births) the sex of last child in a family was negatively correlated with sexes of preceding siblings (probability of male birth decreased with number of preceding brothers) (Thomas Gualtieri, Hicks et al. 1984).

Mitter *et al* reported an excess of unisexual sibships in small sample (451 families) from India (Mitter and Anand 1975).

(Maconochie and Roman 1997) reported that sex ratio was not associated with sexes of preceding siblings in 330,088 sibships from Scotland. (Jacobsen, Moller et al. 1999) used a sample of 815,891 children from Denmark. According to their analysis, neither sex of immediately preceding sibling nor sexes of two or three preceding siblings had significant effect on sex ratio, when the regression model was adjusted for paternal age. (Rodgers and Doughty 2001) analyzed 6,089 families from USA and concluded that sexes of previous siblings had no effect on sex ratio.

Garenne (Garenne 2009) analyzed over 2 million births in Sub-Saharan Africa. There was no correlation between sexes of successive siblings (although, only the correlation between two last births was tested). The sex ratio was dependent on number of preceding boys (positively) and number of preceding girls (negatively), as it was the case in Malinvaud’s sample.

Overall the results of published studies are mixed. Many, but not all of them, report a correlation between sexes of siblings in a family. When the correlation is found, researchers disagree about it’s possible sources, since it can be attributed to both positive Markovian and to Lexian variation. The situation is complicated by a variety of statistical methods employed. Some authors tested pair-wise correlation between sexes of two successive siblings. Others used a regression of probability of male birth on numbers of preceding boys and girls. Methods that enable simultaneous testing of Markovian and Lexian variation were rarely employed. Few studies explicitly tested for an excess of families with particular composition, such as unisex families. Sample size is also an issue, with many studies conducted on only few thousands of families (while others used hundreds of thousands).

Recently large scale demographic data, including birth sequences, became available for analysis from the Demographic and Health Survey (DHS) program. The DHS is an international organization that assists in conducting demographic surveys, mostly in developing countries. It collects and disseminates data on fertility, health, nutrition etc. I used DHS data from about 8 million births (to my knowledge, largest sample used in such studies so far) to test (1) if there are significant deviations from expected frequencies of sibships with various sex compositions, (2) if there is a correlation between sexes of successive siblings, and (3) if this correlation can be attributed to Lexian or Markovian variation.

## Methods

Source data were downloaded from DHS website (dhsprogram.com) in August-September 2015. The demographic surveys were carried out in 1985–2014 mostly in developing countries of Africa, Asia and South America, using standardized questionnaires. Surveys of following types were selected for analysis: Standard DHS, Continuous DHS, Interim DHS, Special DHS. For each survey, the “individual response” dataset (with answers from Individual Woman’s Questionnaire) was downloaded, in Stata format (.dta). The questionnaire included questions about sexes of all children born to respondent woman.

The sequences of siblings sexes were taken from variables b4_01-b4_20. Other variables used in analysis were b11_01-b11_20 (intervals between births), b3_01-b3_20 (child’s date of birth), v011 (mother’s date of birth), b0_01-b0_20 (indicator of plural birth), v201 (total number of children ever born).

Data handling and statistical analysis were done with Stata/MP 13. Logistic regression used sex of each child (except of first child in a sibship, who doesn’t have any preceding sibling) as dependent variable, and sex of previous child – as independent variable. Following observations were excluded from analyses: (1) pairs of successive siblings where at least one of them was from plural birth; (2) observations where recorded value of BTB interval was 9 months or less; (3) sibships with reported total number of children ever born (variable v201) did not match number of children recorded in variables b4_01-b4_20.

Maximum likelihood estimation was performed using the model from (Astolfi and Tentoni 1995). Only sibships with four or less children were analyzed. Briefly, Markovian dependency was modelled with parameters *k_i_*_m_, *k_i_*_f_ (birth order *i* = 2, 3, 4). The probability of *i*’th child being boy equals *k_i_*_m_*p*_1_ when *i-1*’th child was boy and *k_i_*_f_*p*_1_ when *i-1*’th child was girl. The probability of *i*’th child being girl equals 1 - *k_i_*_m_*p*_1_ when *i-1*’th child was boy and 1 - *k_i_*_f_*p*_1_ when *i-1*’th child was girl. Other parameters reflected the shape of probability distribution of *p*_1_ and stopping rules. Markovian variation was excluded by constraining *k_i_*_m_ = *k_i_*_f_ = *k_i_* (*i* = 2, 3, 4); Poisson – by constraining *k_i_*_m_ = *k*_m_, *k_i_*_f_ = *k*_f_ (*i* = 2, 3, 4); Lexian – by constraining 2^nd^, 3^rd^ and 4^th^ central moments of probability distribution to zero. Broyden-Fletcher-Goldfarb-Shanno (BFGS) algorithm was used for likelihood maximization. The code for maximum likelihood estimation was written in Stata, and tested on Astolfi’s data to ensure that it generated same results as in the original publication.

## Results

### Large same-sex sibships were more numerous than expected

Total sample included 7,985,855 children in 2,214,601 sibships (Table 8). A sibship can be characterized, regardless of birth order, by total number of children *n* and number of boys *k.* If sex of each child is determined independently of siblings, then expected frequency of the sibship can be calculated according to binomial law: 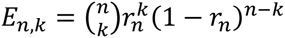, where r_n_ is frequency of boys in all sibships with size *n*. Chi-squared test showed significant difference between expected and observed frequencies (Chi2 = 994, df = 178, p < 0.001). The deviations remained significant when sibships with at least one plural birth were excluded from the analysis (Chi2 = 1071, df = 176, p < 0.001), or when only one frequency of boys *r* for whole sample (instead of specific frequency for each sibship size) was used to calculate expected frequencies (Chi2 = 1700, df = 177, p < 0.001).

Next, chi-squared test was applied to test deviation from expected frequency for each type of sibship (as characterized by *n* and *k*) (Table 1).

**Table 1.** Odds ratios demonstrating relative abundance of sibships with size *n* and number of boys *k*. Odds ratio above 1 indicates an excess of sibships with given composition, relative to expected frequency. *p-value < 0.05. For 20 out of 63 chi-square tests listed in the table the p-value was below 0.05. Sibships with n > 10 are not shown. Sibships with at least one plural birth were excluded.

| $k$ | $n$ | | | | | | | | |
| --- | --- | --- | --- | --- | --- | --- | --- | --- | --- |
|  | 2 | 3 | 4 | 5 | 6 | 7 | 8 | 9 | 10 |
| 0 | 0.94* | 0.94* | 0.92* | 0.95* | 0.96 | 1.04 | 1.21* | 1.17 | 1.36 |
| 1 | 1.1* | 1.01* | 0.99 | 1.01 | 1.01 | 1.04 | 1.05 | 1.18* | 1.14 |
| 2 | 0.94* | 1.05* | 1.05* | 1.01 | 1.01 | 1.01 | 1.02 | 1.04 | 1.07 |
| 3 |  | 0.93* | 0.98* | 0.98* | 0.99 | 0.98 | 0.99 | 0.99 | 1 |
| 4 |  |  | 0.96* | 1.01 | 0.98* | 0.99 | 0.99 | 0.97 | 0.99 |
| 5 |  |  |  | 1 | 1.02 | 0.99 | 0.97 | 0.99 | 0.99 |
| 6 |  |  |  |  | 1.04 | 1.04 | 1.02 | 0.98 | 0.98 |
| 7 |  |  |  |  |  | 1.13* | 1.05 | 1.04 | 0.97 |
| 8 |  |  |  |  |  |  | 1.29* | 1.15* | 1.06 |
| 9 |  |  |  |  |  |  |  | 0.98 | 1.26* |
| 10 |  |  |  |  |  |  |  |  | 1.59 |

The deviations were significant for many types of sibship. Notably, sibships with many boys were in excess among large families. For instance, among 24,999 sibships with exactly ten children (and no plural births) 43 were all-boys, while expected number was only 27 (OR = 1.59, p = 0.07). Similarly, among large sibships number of those composed mostly with girls tended to be higher than expected. At the same time among small sibships (<5 children) mixed-sex types were more common. For example, among sibships with only 2 children those with one boy and one girl were overrepresented (OR = 1.10, p < 0.001).

### Sex ratio depended on the sex of preceding sibling

The dependence between sexes of successive siblings was tested using logistic regression (boy coded as “0” and girl as “1”). The sex of preceding sibling had small but significant positive effect on sex of a child (coef = 0.016, p < 0.001, ~0.4%^3^ change of sex ratio). Assuming that this effect could depend on sibship size and birth order of a child, the regression was repeated for different sibship sizes *n* and birth orders *i* (Table 2).

**Table 2.** Correlations between sexes of successive siblings, by sibship size *n* and birth order *i.* Regression coefficients from logit model are presented (sex of *i*’th child as dependent variable, sex of *i-1*’th child – independent variable). Sibships with more than ten children are not shown. *p-value < 0.05.

| $i$ | $n$ | | | | | | | | |
| --- | --- | --- | --- | --- | --- | --- | --- | --- | --- |
|  | 2 | 3 | 4 | 5 | 6 | 7 | 8 | 9 | 10 |
| 2 | -0.186* | 0.011 | 0.069* | 0.103* | 0.089* | 0.074* | 0.106* | 0.089* | 0.039 |
| 3 |  | -0.13* | -0.004 | 0.077* | 0.089* | 0.091* | 0.089* | 0.1* | 0.179* |
| 4 |  |  | -0.088* | 0.011 | 0.091* | 0.086* | 0.093* | 0.105* | 0.119* |
| 5 |  |  |  | -0.039* | 0 | 0.1* | 0.089* | 0.114* | 0.101* |
| 6 |  |  |  |  | -0.037* | 0.034* | 0.085* | 0.08* | 0.066* |
| 7 |  |  |  |  |  | -0.018 | 0.07* | 0.072* | 0.068* |
| 8 |  |  |  |  |  |  | 0.005 | 0.068* | 0.041 |
| 9 |  |  |  |  |  |  |  | -0.002 | 0.046 |
| 10 |  |  |  |  |  |  |  |  | 0.064* |

When dependent variable was the sex of last child in a sibship (that is, *i* = *n*), the regression coefficients were mostly negative. Meanwhile, for most other combinations of *n* and *i* the coefficients were positive and significant. When the analysis (for all sibship sizes *n* combined) was performed excluding last child of each sibship (i.e. when *i* ≠ *n,* for all *n*), the regression coefficient was 0.067 (p < 0.001). Conversely, when only last children were included (*i* = *n*, for all *n*), the regression coefficient was -0.098 (p < 0.001).

For confirmation of the results obtained in pooled sample the regression was performed in each dataset from DHS website (one dataset per survey), and then random effects meta-analysis was applied to pool estimates of regression coefficients. When all children were included (regardless of birth order) the result was not significant (coef = 0.017, p = < 0.001; I^2^ = 46%, p < 0.001). When last child of each sibship was excluded, the effect of preceding sibling’s sex was positive and significant (coef = 0.063, p < 0.001; I^2^ = 57.7%, p < 0.001). When only last children were included the effect of previous sibling’s sex was negative and significant (coef = -0.084, p < 0.001; I^2^ = 79.6%, p < 0.001).

Why does the sign of this correlation depend on whether “dependent” child is last or not last in a family? Is it possible that sex of preceding sibling affects sex of a child via two different mechanisms, depending on whether the next sibling is a last child in the family or not? The negative correlation for last child may be explained by deliberate family planning, namely parents’ willingness to have a family with mixed sex composition (“balance preference”, discussed by (Gini 1951) and others). Such preference may dictate specific stopping behavior. For instance, if a couple have several daughters they may continue procreation until having a son, and then stop. A couple having a boy and a girl may decide to stop the procreation, while a family with two girls may wish to have one more child in a hope that it will be a boy.

However, the last child recorded in DHS data is not necessary the last child in the family. The respondent woman may continue the procreation after interview date (i.e. the family could be incomplete). The proportion of children who are last in complete families should increase with the interval between date of last birth and date of interview. If the negative correlation between sexes of two last siblings is caused by parents’ stopping behavior, the correlation should be stronger when proportion of complete families is higher. Indeed, in the logit model including the time since last birth as a covariate, the effect of interaction term was negative and significant (Table 3, column *3*). The negative sign suggests that coefficient’s absolute value is increasing when the time since last birth increases. For instance, when last birth was less than 6 months before interview (bottom 10% of distribution of interval values) the effect of previous sibling’s sex was not significant (coef = -0.018, p = 0.088). When last birth was more than 167 months before interview (top 10%) the negative correlation was highly significant (coef = -0.25, p < 0.001), presumably because the proportion of complete families in this subsample was high.

**Table 3.**
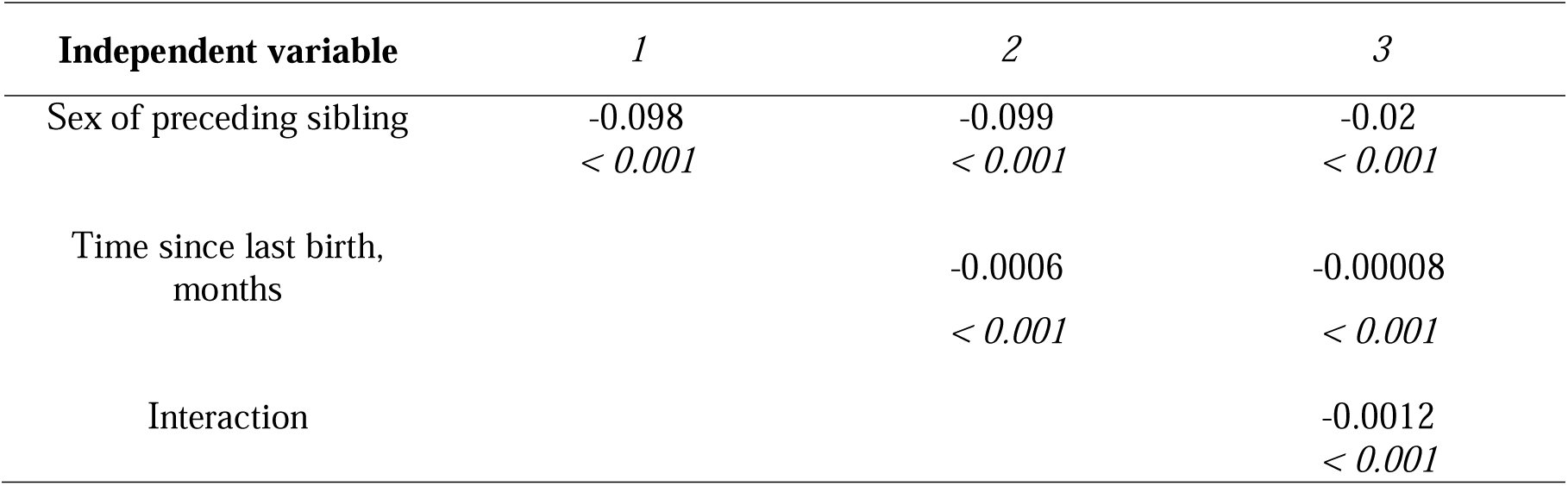
Effects of preceding sibling’s sex and time since last birth on sex of last sibling in a sibship. Logistic regression coefficients. p-values in italic.

| Independent variable | 1 | 2 | 3 |
| --- | --- | --- | --- |
| Sex of preceding sibling | -0.098<br>*< 0.001* | -0.099<br>*< 0.001* | -0.02<br>*< 0.001* |
| Time since last birth,<br>months |  | -0.0006<br>*< 0.001* | -0.00008<br>*< 0.001* |
| Interaction |  |  | -0.0012<br>*< 0.001* |

Alternatively to deliberate family planning, some biological factors behind the correlation between sexes may change their direction from positive to negative, and such a change could be accompanied by the end of procreation.

### Was the correlation caused by Lexian or Markovian variation?

If some parents have a tendency to produce children of particular sex, then a correlation should be observed between sexes of siblings regardless their birth order, not only between successive siblings (Edwards 1961). If the association between successive siblings has major role, then little or no correlation should be observed between sexes of siblings separated by other births. In DHS data the correlation was strong between successive siblings, less strong but still significant– between siblings separated by one birth, and negligible when siblings were separated by two or more births (Table 4, column *2*).

**Table 4.** Correlations between sexes of siblings separated by various numbers of births. Logistic regression coefficients. Sibships with more than 10 children are not shown.

| Number<br>of births<br>separating<br>the two<br>siblings | 1<br>All siblings |  |  | 2<br>Excluding last children |  |  | 3<br>Only last children |  |  |
| --- | --- | --- | --- | --- | --- | --- | --- | --- | --- |
|  | *Coef* | *P* | *N* | *coef* | *p* | *N* | *coef* | *p* | *N* |
| 0 | 0.016 | <0.001 | 5497715 | 0.067 | <0.001 | 3781932 | -0.098 | <0.001 | 1715783 |
| 1 | -0.011 | <0.001 | 3766133 | 0.021 | <0.001 | 2519460 | -0.077 | <0.001 | 1246673 |
| 2 | -0.019 | <0.001 | 2524914 | 0.003 | 0.33 | 1648143 | -0.059 | <0.001 | 876771 |
| 3 | -0.015 | <0.001 | 1653095 | 0.004 | 0.27 | 1046955 | -0.049 | <0.001 | 606140 |
| 4 | -0.013 | <0.001 | 1051987 | -0.008 | 0.13 | 638499 | -0.022 | <0.001 | 413488 |
| 5 | -0.014 | 0.004 | 643115 | -0.006 | 0.37 | 370077 | -0.026 | <0.001 | 273038 |
| 6 | -0.003 | 0.70 | 373785 | 0.004 | 0.63 | 201077 | -0.01 | 0.28 | 172708 |
| 7 | 0.001 | 0.93 | 203653 | -0.006 | 0.63 | 101588 | 0.008 | 0.54 | 102065 |
| 8 | -0.001 | 0.96 | 103270 | 0.004 | 0.83 | 46867 | -0.004 | 0.80 | 56403 |
| 9 | 0.004 | 0.82 | 47840 | -0.015 | 0.59 | 20087 | 0.018 | 0.45 | 27753 |
| 10 | -0.018 | 0.52 | 20606 | -0.037 | 0.42 | 7770 | -0.007 | 0.85 | 12836 |

When only last children were concerned (i.e., last sibling’s sex was dependent variable) the negative coefficient remained highly significant even when siblings were separated by up to five births (Table 4, column 3). This negative correlation with sexes of several preceding siblings is not surprising if the last child’s sex is influenced by parents’ balance preference (parents take into consideration the sexes of several existing children).

When all children, regardless of birth order, were included in the model, the coefficient was positive and significant for successive births, but it was negative and significant when siblings were separated by 1–5 births (Table 4, column 1). Apparently, this result is caused by combination of two effects: positive correlations in most successive pairs, and negative correlation between sex of last child and sexes of several siblings preceding the last one.

More formal way to test the presence of Markovian dependency is to use a statistical model where impacts of Markovian, Lexian and Poisson variations are represented simultaneously by different sets of parameters (see Methods). Likelihood ratios can be used to test significance of particular parameters for model fit. Four models were fit to DHS data: a model with all parameters being estimated (full model), with Markovian variation excluded (Model 1), with Poisson variation excluded (Model 2) and with both Markovian and Lexian variation excluded (Model 3). Only sibships with four or less siblings were considered (71% of all sibships). The influences of all three sources of variation were significant: Markovian (LR = 82, df = 3, p < 0.001), Poisson (LR = 31, df = 4, p < 0.001) and Lexian (LR = 910, df = 3, p < 0.001). If Lexian variation is mediated by beta-distribution of probability of male birth, then evaluated distribution parameters are *a* = 64.1 and *b* = 60.3. The variance of this distribution is 0.00196, which is close to Edwards’ estimate of 0.0025 (Edwards 1958).

### Effects of birth-to-birth interval on the correlation between sexes

If previous sibling’s sex exerts it’s effect by leaving some kind of temporary chemical mark in mother’s organism, then this effect could be stronger for shorter intervals between births (Edwards 1961). The interaction between birth-to-birth (BTB) interval and sex of previous sibling was tested using logistic regression (Table 5).

**Table 5.** Interaction of preceding sibling’s sex and birth-to-birth (BTB) interval. Logistic regression coefficients are shown (p-values in italic).

| Independent variable | Excluding last children |  |  | Only last children |  |  |
| --- | --- | --- | --- | --- | --- | --- |
|  | 1 | 2 | 3 | 4 | 5 | 6 |
| Sex of preceding sibling | 0.067<br>*<0.001* | 0.067<br>*<0.001* | 0.095<br>*<0.001* | -0.098<br>*<0.001* | -0.098<br>*<0.001* | -0.096<br>*<0.001* |
| BTB interval, months |  | 0.0004<br>*<0.001* | 0.0009<br>*<0.001* |  | 0.0007<br>*<0.001* | 0.0007<br>*<0.001* |
| Interaction term |  |  | -0.0009<br>*<0.001* |  |  | 0.00005<br>*0.6* |

The effect of BTB interval alone was significant, regardless of whether last children were excluded or not: every additional month since last birth increased chances to have a girl by approximately 0.01%. The interaction term was significant only in the model excluding last children (coef = -0.0009, p < 0.001). The negative sign of the coefficient suggests that the effect of preceding sibling’s sex is vanishing with time. This interaction can be seen on Figure 1 (the value of regression coefficient is decreasing from 9 to 40–45 months of BTB).

**Figure 1.**
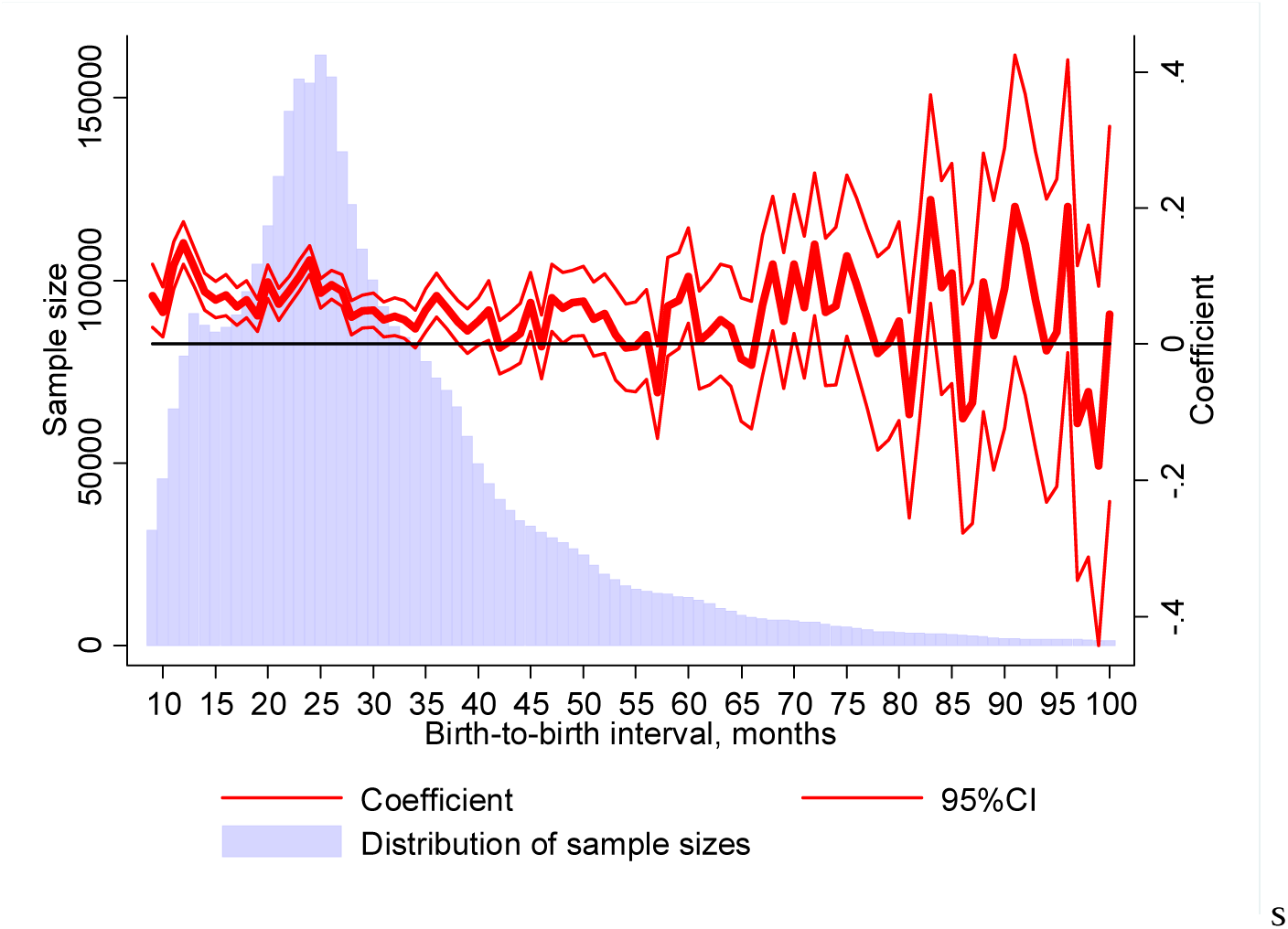
Logistic regression coefficients (effect of preceding sibling’s sex) by BTB intervals, with last child of each sibship excluded from analysis. Distribution of sample sizes over BTB values is shown in the background.

When the analysis was limited to last children, the interaction was not significant (coef = - 0.00003, p = 0.8), Table 5, Figure 2.

**Figure 2.**
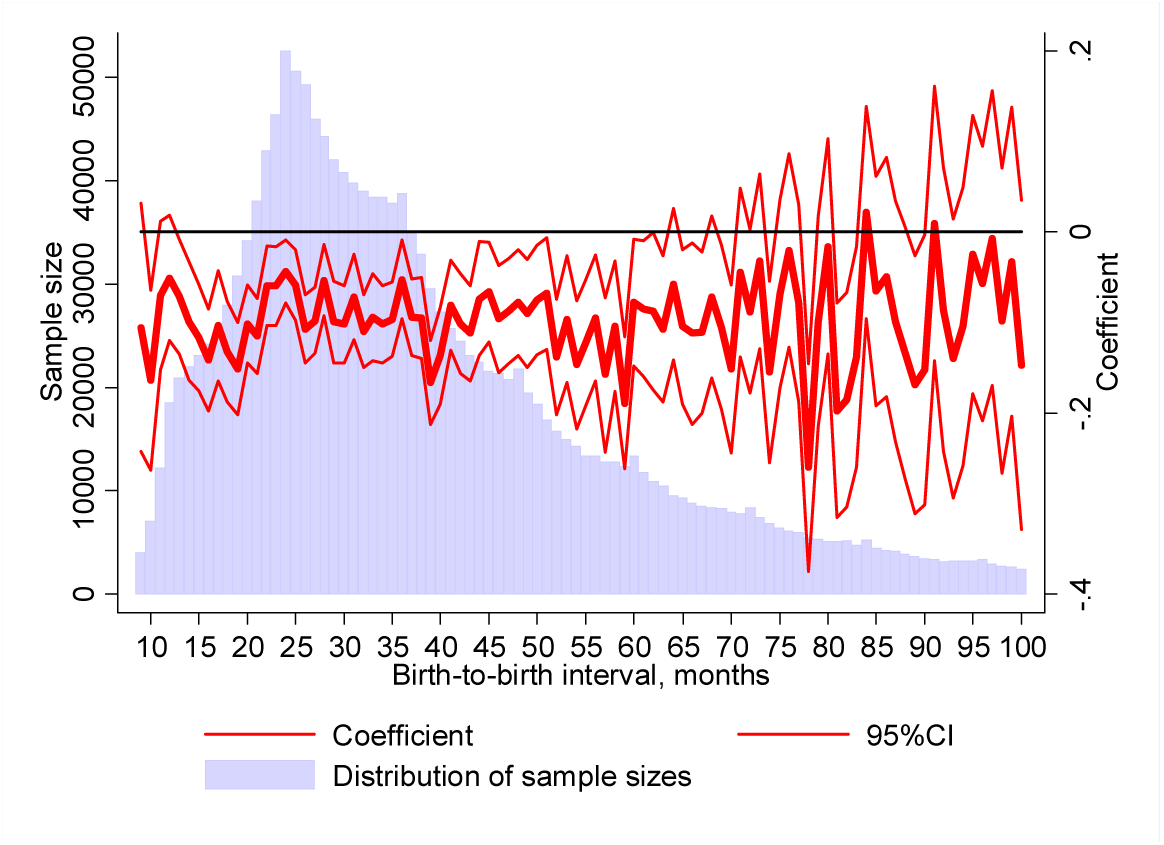
The effect of preceding sibling’s sex doesn’t change substantially with BTB interval, when only last children are considered. Logistic regression coefficients (effect of preceding sibling’s sex) by BTB intervals. Distribution of sample sizes over interval values is shown in the background.

Edwards (Edwards 1961) suggested also that the correlation between successive siblings’ sexes could be mediated by secondary adjustment of sex ratio, namely by lower chances of fetus survival when it’s sex is opposite to preceding sibling. In this case the BTB interval could be longer when successive siblings were of unlike sex.

BTB interval between same sex births was significantly shorter than between opposite sex births when excluding last children (linear regression, pair of same sex births coded as “1”, opposite sex − “0”, coef = -0.135, p < 0.001). Results of nonparametric rank sum test were significant as well (p < 0.001).

(Greenberg and White 1967) noted that such analysis of intervals should control for birth order or family size (because if their data the interval decreased with family size when birth order was constant). When sibship size and birth order were included into regression model as covariates, the interval still was significantly longer for opposite sex pairs (coef = -0.085, p < 0.001). Additionally, the analysis was repeated for each combination of sibship size *n* and birth order *i* (Table 6).

**Table 6.** Differences between BTB intervals for siblings of same and opposite sex. Linear regression coefficients are shown (the value corresponds to difference of BTB between pairs of opposite sex siblings and pairs of same sex siblings). * p < 0.05. Sibships with more than 10 children are not shown.

| *i* | *n* |  |  |  |  |  |  |  |
| --- | --- | --- | --- | --- | --- | --- | --- | --- |
|  | 3 | 4 | 5 | 6 | 7 | 8 | 9 | 10 |
| 2 | 0.337* | 0.03 | -0.142 | -0.008 | -0.263* | -0.124 | -0.167 | -0.047 |
| 3 |  | 0.257* | -0.055 | -0.127 | -0.309* | -0.331* | -0.071 | -0.284* |
| 4 |  |  | 0.019 | -0.072 | -0.11 | -0.339* | -0.137 | -0.108 |
| 5 |  |  |  | -0.108 | -0.268* | -0.333* | -0.164 | -0.24 |
| 6 |  |  |  |  | -0.308* | -0.015 | -0.278* | -0.367* |
| 7 |  |  |  |  |  | -0.167 | -0.405* | -0.115 |
| 8 |  |  |  |  |  |  | -0.176 | -0.273 |
| 9 |  |  |  |  |  |  |  | -0.372* |

The difference of BTB intervals was significant for multiple, but not for all, combinations of *n* and *i.* Notably, all coefficients were negative for sibships with six or more children.

## Additional datasets

Detailed information about siblings sexes and birth order was presented in several published papers ((Renkonen, Makela et al. 1961, Greenberg and White 1967, Maconochie and Roman 1997, Jacobsen, Moller et al. 1999, Rodgers and Doughty 2001)). Logistic regression was applied to these datasets to investigate the reproducibility of results obtained in DHS data (Table 7).

**Table 7.** Logistic regressions of child’s sex on preceding siblings’ sex in datasets from published papers.

| **Publication** |  | **All observations** | **Excluding last child** | **Limited to last child** |
| --- | --- | --- | --- | --- |
| (Renkonen, Makela et al. 1961) (as presented in (Edwards 1962))<sup>1</sup> |  |  |  |  |
| *All data* | *coef* | 0.053 | 0.104 | -0.017 |
|  | *p* | < 0.001 | < 0.001 | 0.33 |
|  | *N* | 123445 | 71577 | 51868 |
| *< 5 siblings* | *coef* | 0.037 | 0.159 | -0.05 |
|  | *p* | 0.021 | < 0.001 | 0.018 |
|  | *N* | 62797 | 26091 | 36706 |
| (Greenberg and White 1967)<sup>2</sup> |  |  |  |  |
| *All data* | *coef* | -0.008 | 0.005 | -0.044 |
|  | *p* | 0.195 | 0.46 | < 0.001 |
|  | *N* | 438788 | 322399 | 116389 |
| *< 7 siblings* | *coef* | -0.024 | 0.004 | -0.073 |
|  | *p* | 0.005 | 0.72 | < 0.001 |
|  | *N* | 220154 | 140204 | 79950 |
| (Maconochie and Roman 1997) |  |  |  |  |
|  | *coef* | -0.004 | 0.512 | -0.141 |
|  | *p* | 0.7 | < 0.001 | < 0.001 |
|  | *N* | 218028 | 46159 | 171869 |
| (Jacobsen, Moller et al. 1999)<sup>3</sup> |  |  |  |  |
|  | *coef* |  | 0.315 |  |
|  | *p* |  | < 0.001 |  |
|  | *N* |  | 45757 |  |
| (Rodgers and Doughty 2001)<sup>4</sup> |  |  |  |  |
| *All data* | *coef* | -0.065 | 0.076 | -0.14 |
|  | *p* | 0.196 | 0.372 | 0.023 |
|  | *N* | 6435 | 2227 | 4208 |
| *< 4 siblings* | *coef* | -0.153 | -0.036 | -0.195 |
|  | *p* | 0.007 | 0.74 | 0.003 |
|  | *N* | 5046 | 1301 | 3745 |

<sup>1</sup> in original report families with five or more children were pooled together. Thus, the analysis limited to last children was informative only for sibships with four siblings or less. <sup>2</sup> in original report families with seven or more children were pooled. Thus, the analysis limited to last children was informative only for sibships with six siblings or less. <sup>3</sup> the data were taken from Table V of original report, that doesn't lists last children. Thus, only the analysis excluding last children (and only in sibships with three or more siblings) was possible. <sup>4</sup> in original report families with four or more children were pooled. Thus, the analysis limited to last children was informative only for sibships with three siblings or less.

In data from Renkonen *et al* (Renkonen, Makela et al. 1961) the effect of previous sibling’s sex was positive when last children were excluded (coef = 0.158, p< 0.001) and negative when only last children were included (coef = -0.049, p = 0.018), as in the DHS data. In Greenberg’s data (Greenberg and White 1967) positive correlation, after exclusion of last children, was not significant, while there was strong negative correlation between sexes of last two siblings in a family. Similar situation was observed in Rodgers data (Rodgers and Doughty 2001) (excluding last children - not significant effect; including only last children, coef = -0.19, p = 0.003). In Maconochie’s data (Maconochie and Roman 1997) both kinds of correlation were significant. In Jacobsen’s data (Jacobsen, Moller et al. 1999) the correlation was positive and significant in the model excluding last children (coef = 0.078, p < 0.001).

Thus, positive correlation between sexes of successive siblings (excluding last children) was found in three out of five additional datasets. It should be noted, however, that Jacobsen and Rodgers do not report statistics of birth sequences after excluding plural births. Thus the positive values of coefficients could be overestimated. One should note also that unlike other datasets, Renkonen data don’t include children from a previous marriage of each woman.

## Discussion

Current results demonstrate an excess of large sibships with high (many boys) and with low (many girls) sex ratios, relative to binomial expectation. The overrepresentation of such sibships was accompanied by a correlation between the sexes of siblings. Notably, positive correlation was observed between the sexes of successive siblings and siblings separated by one birth, but not for those separated by two or more births. This effect is consistent with the hypothesis of Markovian variation, but not with Lexian one. However, formal modelling indicated that both Markovian and Lexian variation were present.

Makela (Makela 1963) noted that in the presence of Lexian variation, a lack of correlation for siblings separated by several births could be observed due to other factors, for example an immunization against male fetal antigens. But in this case it is not clear why such factors do not make the correlation between successive siblings negative as well.

Additional evidence for Markovian variation comes from the analysis of birth-to-birth intervals. Namely, the correlation between sexes was stronger when births were separated by shorter intervals.

Several published studies reported an absence of Markovian variation, but a negative report doesn’t necessarily mean that the correlation is absent. Perhaps it is absent in a subsample selected for the testing, or can’t be detected using specific methods employed. For example, Garenne (Garenne 2009) performed a test of association between the sexes of successive siblings and found it insignificant. However, this test was conducted not on all children in the sample. Instead, from each sibship only two siblings were selected: the last one and the second to last in birth sequence. Moreover, the analysis was restricted to women at least 35 years old.

Jacobsen et al (Jacobsen, Moller et al. 1999) and Maconochie and Roman (Maconochie and Roman 1997) reported that the correlation between sexes was not significant, but no distinction was made between last birth in a sibship and all other births. When logistic regression was applied only to non-last births, significant positive association was seen between successive siblings (Table 7).

Regarding the sample used in (Greenberg and White 1967) sample, there seems to be no positive Markovian correlation even when the non-last births were tested separately. The lack of correlation in this sample might be explained by peculiarities of the source data. The birth records were from archives of the Genealogical Society of the Church of Jesus Christ of Latter Day Saints (also known as Mormons) at Salt Lake City, Utah, USA. The fathers of these families were born on or after the year 1800. Some Mormons practiced polygamy during that period, at least until 1910 (Wikipedia 2015). If a man had more than one wife, the progeny of each wife was recorded as a separate sibship. Also, if a woman had children with more than one man, the offspring of each man was recorded as a separate sibship. These peculiarities could potentially hinder Markovian correlation. Moreover, some of the children were born not in USA, but in Norway, Sweden, Denmark and Holland, i.e. before their parents moved to United States. These circumstances could reduce the quality of the data and further contribute to the discrepancy between the study findings and results from other studies. Notably, the authors re-analyzed Renkonen’s data (Renkonen, Makela et al. 1961) and confirmed that the correlation was present in that sample.

It is unknown what biological mechanisms could underlie positive Markovian variation. Gini (Gini 1951) stated that “[the] hypothesis may be considered excluded by the mechanism of sex determination in a male heterogametic species like the humans”. Other researchers also noted that there is no plausible mechanism for the influence of one siblings’ sex on the next one (Ben-Porath and Welch 1976, James 2000).

However, the casual Markovian dependence is not biologically impossible. At least two potential mechanisms can be proposed. First is an influence of testosterone. Testosterone is produced by male fetuses (Conte and Grumbach 2011), and the male fetuses have higher testosterone levels in cord blood (Maccoby, Doering et al. 1979, Nagamani, McDonough et al. 1979, Herruzo, Mozas et al. 1993), compared to females. Maternal testosterone correlates positively with offspring sex ratio in cows (Grant and Irwin 2005) and field voles (Helle, Laaksonen et al. 2008), i.e. it increases the proportion of males. There is a lot of indirect evidence that testosterone can increase sex ratio in humans too (James 2011). The mechanism of these correlations is unknown, but the influence of testosterone could be long lasting: an exposure of mother to fetal testosterone during a pregnancy may trigger physiological changes that will influence the sex ratio of the next offspring.

Another potential mechanism of Markovian dependency could be based on fetal microchimerism. During pregnancy fetal cells penetrate into mother’s organism and remain there for years. These cells migrate into heart, skin, thyroid and adrenal glands, liver, kidney, lung, and spleen (Bayes-Genis, Roura et al.). Presumably, they may migrate into reproductive system too. They are able to differentiate into myocardial cells (Bayes-Genis, Roura et al.) and probably into other cell types (Khosrotehrani and Bianchi 2005). In principle, such cells, remaining in the reproductive system after previous pregnancy, may induce an adjustment of sex ratio by influencing viscosity of cervical mucus or embryo mortality. Alternatively, fetal cells may alter activity of adrenals (and levels of cortisol), pituitary (and levels of ACTH), ovaries (and levels of sex steroids) and other glands related to sex ratio adjustment.

A study on Mongolian gerbils demonstrated that females, developed between male embryos while *in utero,* had higher sex ratio in their offspring later on (Clark and Galef 1995). This effect in gerbils could be related to an exposure of female fetus to testosterone from adjacent male fetuses, or an acquisition of cells from male fetuses. Perhaps this effect is based on same mechanism as the Markovian correlation in humans.

One study reported that concentration of fat in human milk depends on child’s sex (Fujita, Roth et al. 2012). The authors noted also “near-significant negative influence of the number of sons living at home on milk fat concentrations”. Biological mechanism of these effects is unknown, but it is likely that sex of the child causes persistent alterations in mother’s physiology, that can be related to both milk production and adjustment of sex ratio.

The positive markov correlation may have adaptive evolutionary value. There is a famous model of sex ratio adjustmet in the course of evolution, known as Fisher’s principle. According to that model, mutations, which enhance production of children of either sex, should be favored by natural selection if that sex is rare in the population. (It explains why the sex ratio should be close to 1:1 under normal circumstances.) When a mutation alters sex ratio in an offspring, this alteration can be intensified by positive markov dependency. If the change of sex ratio is adaptive (say, a mutation increases proportion of sons when there are too many women in the population) the markov dependency raises its value. Thus, a mechanism underlying Markov dependency would be under positive selection too. Surely, not every trait is under effective selection, and this dependency could be a kind of “spandrel” (Gould and Lewontin 1979), a by-product of unknown physiological mechanisms.

**Future directions.** Among many published studies of sex ratios only few present raw data. Most papers contain only a summary of data, that isn’t sufficient for independent analysis. Presenting full data is essential for replication. In case of sex sequences the data can be made quite compact: it is enough to indicate birth sequence (e.g., Boy-Girl-Boy) and number of sibships of each type. The data used in current paper are available on DHS website (after an approval from DHS).

Biological mechanisms of Markovian variation could be investigated using model organisms. To the best of my knowledge, among animals the Markovian dependence (controlling for Lexian variation) was explicitly tested only in dairy cattle (Astolfi and Tentoni 1995). The variation was not significant, although authors noted weak effect in expected direction: “after a male calf the probability of a male birth was slightly higher than the probability of a female birth”. It would be interesting to see if Markovian variation is present in mice, rats or other species amenable to experimentation.

## Acknowledgements

I’m grateful to all participants of DHS program for collection of the data and making them available for analysis. The possible evolutionary contingencies of Markov correlation were suggested by Dr Ivan Kuzin (Mosow State University, Department of Philosophy).Supplementary information

---

**Table 8.**
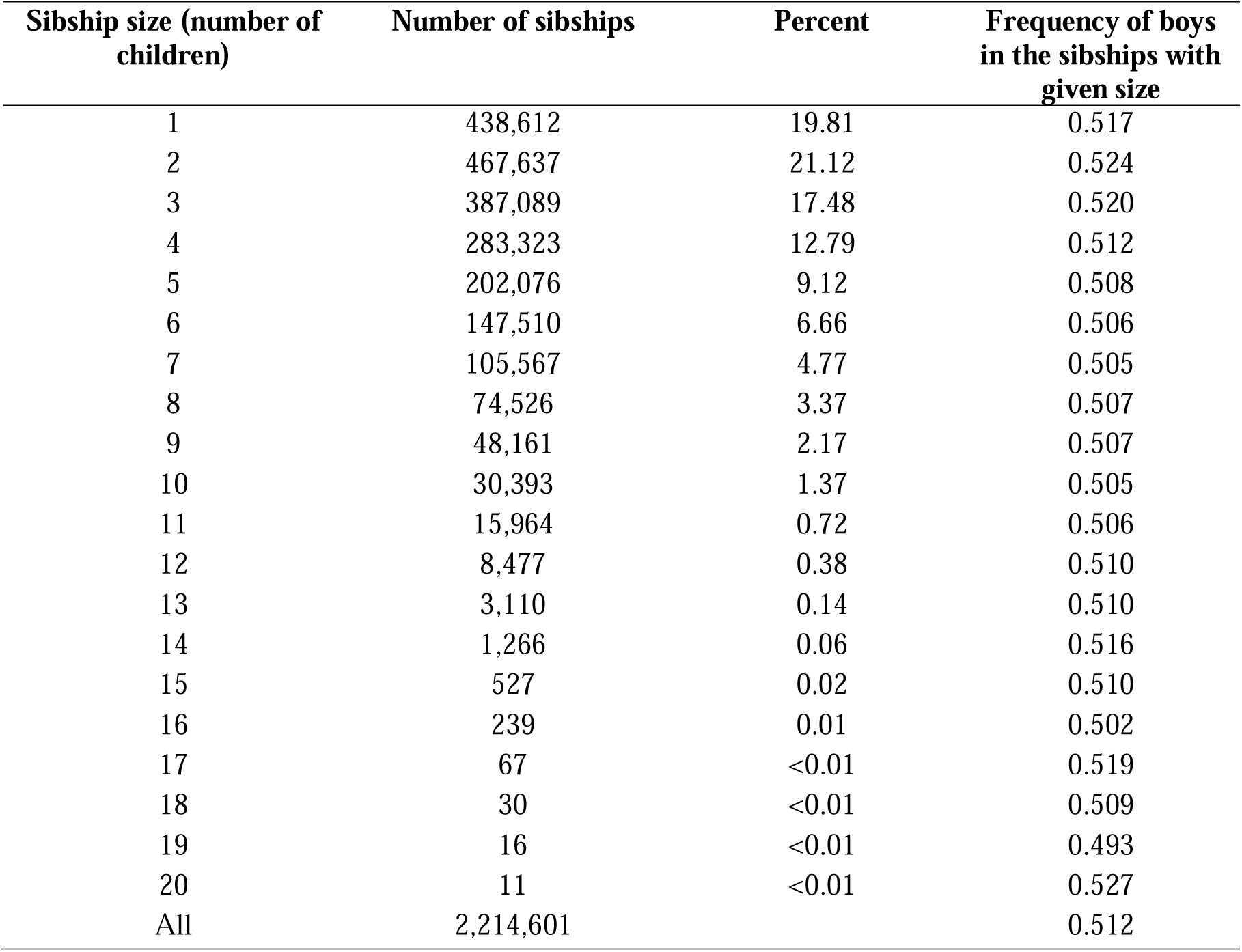
Summary statistics

| Sibship size (number of children) | Number of sibships | Percent | Frequency of boys in the sibships with given size |
| --- | --- | --- | --- |
| 1 | 438,612 | 19.81 | 0.517 |
| 2 | 467,637 | 21.12 | 0.524 |
| 3 | 387,089 | 17.48 | 0.520 |
| 4 | 283,323 | 12.79 | 0.512 |
| 5 | 202,076 | 9.12 | 0.508 |
| 6 | 147,510 | 6.66 | 0.506 |
| 7 | 105,567 | 4.77 | 0.505 |
| 8 | 74,526 | 3.37 | 0.507 |
| 9 | 48,161 | 2.17 | 0.507 |
| 10 | 30,393 | 1.37 | 0.505 |
| 11 | 15,964 | 0.72 | 0.506 |
| 12 | 8,477 | 0.38 | 0.510 |
| 13 | 3,110 | 0.14 | 0.510 |
| 14 | 1,266 | 0.06 | 0.516 |
| 15 | 527 | 0.02 | 0.510 |
| 16 | 239 | 0.01 | 0.502 |
| 17 | 67 | <0.01 | 0.519 |
| 18 | 30 | <0.01 | 0.509 |
| 19 | 16 | <0.01 | 0.493 |
| 20 | 11 | <0.01 | 0.527 |
| All | 2,214,601 |  | 0.512 |

## Footnotes

1 The probability of male birth should be distinguished from sex ratio, which is number of boys per 100 girls. But if the probability is calculated as a frequency of boys, then higher probability implies higher sex ratio and vice versa.

2 Other aspects of sex ratio variation were studied as early as in 1710, see James, W. H. (2000). “The variation of the probability of a son within and across couples.” Human Reproduction **15**(5): 1184–1188.

3 As a rule of thumb in these models, logistic regression coefficient of 0.1 corresponds to 2.5% increase of probability that the *i*’th child has same sex as preceding sibling.

## References

Astolfi, P. and S. Tentoni (1995). “Sources of variation of the cattle secondary sex ratio.” Genetics Selection Evolution 27(1): 3–14.

Bayes-Genis, A., S. Roura, C. Prat-Vidal, J. Farre, C. Soler-Botija, A. B. de Luna and J. Cinca “Chimerism and microchimerism of the human heart: evidence for cardiac regeneration.” Nat Clin Pract Cardiovasc Med.

Beilharz, R. G. (1963). “A factorial analysis of sex-ratio data. A comment on two papers by Edwards.” Ann Hum Genet 26: 355–358.

Ben-Porath, Y. and F. Welch (1976). “Do Sex Preferences Really Matter?” The Quarterly Journal of Economics 90(2): 285–307.

Bernstein, M. E. (1952). “Studies in the human sex ratio: 2. The proportion of unisexual sibships.” Human Biology 24(1): 35–43.

Clark, M. M., and B. G. Galef, Jr. (1995). “A gerbil dam’s fetal intrauterine position affects the sex ratios of litters she gestates.” Physiol Behav 57(2): 297–299.

Conte, F. A., and M. M. Grumbach (2011). Chapter 14. Disorders of Sex Determination and Differentiation. Greenspan’s Basic & Clinical Endocrinology, 9e. D. G. Gardner and D. Shoback. New York, NY, The McGraw-Hill Companies.

Edwards, A. W. (1958). “An analysis of Geissler’s data on the human sex ratio.” Annals of human genetics 23(1): 6–15.

Edwards, A. W. (1961). “A factorial analysis of sex ratio data.” Ann Hum Genet 25: 117–121.

Edwards, A. W. (1962). “A factorial analysis of sex-ratio data: a correction to the article in Vol. 25, 117, including some new data and a note on computation.” Ann Hum Genet 25: 343–346.

Edwards, A. W. (1966). “Sex-ratio data analysed independently of family limitation.” Ann Hum Genet 29(4): 337–347.

Fujita, M., E. Roth, Y. J. Lo, C. Hurst, J. Vollner and A. Kendell (2012). “In poor families, mothers’ milk is richer for daughters than sons: a test of Trivers-Willard hypothesis in agropastoral settlements in Northern Kenya.” Am J Phys Anthropol 149(1): 52–59.

Garenne, M. (2009). “Sex ratio at birth and family composition in sub-saharan africa: Inter-couple variations.” Journal of Biosocial Science 41(3): 399–407.

Geissler, A. (1889). “Beitrage zur Frage des Geschlechterverhaltnisses der Geborenen.” Zeitschrift des Königlich-Sächsischen Statistischen Bureaus 35: 1–24.

Gini, C. (1951). “Combinations and sequences of sexes in human families and mammal litters.” Human Heredity 2(3): 220–244.

Gould, S. J., and R. C. Lewontin (1979). “The Spandrels of San Marco and the Panglossian Paradigm: A Critique of the Adaptationist Programme. “ Proceedings of the Royal Society of London B: Biological Sciences 205(1161): 581–598.

Grant, V. J., and R. J. Irwin (2005). “Follicular fluid steroid levels and subsequent sex of bovine embryos. “ Journal of Experimental Zoology Part A: Comparative Experimental Biology 303 (12) : 1120–1125.

Greenberg, R. A., and C. White (1967). “The sexes of consecutive sibs in human sibships.” Human Biology 39(4): 374–404.

Helle, S., T. Laaksonen, A. Adamsson, J. Paranko and O. Huitu (2008). “Female field voles with high testosterone and glucose levels produce male-biased litters.” Animal Behaviour 75(3): 1031–1039.

Herruzo, A. J., J. Mozas, J. L. Alarcon, J. M. Lopez, R. Molina, L. Molto and J. Martos (1993). “Sex differences in serum hormone levels in umbilical vein blood.” Int J Gynaecol Obstet 41(1): 37–41.

Jacobsen, R., H. Moller and A. Mouritsen (1999). “Natural variation in the human sex ratio.” Hum Reprod 14(12): 3120–3125.

James, W. H. (1975). “Sex ratio and the sex composition of the existing sibs.” Annals of Human Genetics 38(3): 371–378.

James, W. H. (2000). “The variation of the probability of a son within and across couples.” Human Reproduction 15(5): 1184–1188.

James, W. H. (2011). “The categories of evidence relating to the hypothesis that mammalian sex ratios at birth are causally related to the hormone concentrations of both parents around the time of conception.” Journal of Biosocial Science 43(2): 167–184.

Khosrotehrani, K. and D. W. Bianchi (2005). “Multi-lineage potential of fetal cells in maternal tissue: a legacy in reverse.” Journal of Cell Science 118(8): 1559–1563.

Maccoby, E. E., C. H. Doering, C. N. Jacklin and H. Kraemer (1979). “Concentrations of sex hormones in umbilical-cord blood: their relation to sex and birth order of infants.” Child Dev 50(3): 632–642.

Maconochie, N. and E. Roman (1997). “Sex ratios: are there natural variations within the human population?” Br J Obstet Gynaecol 104(9): 1050–1053.

Makela, O. (1963). “Analysis of some sex ratio data.” Ann Hum Genet 26: 333–334.

Malinvaud, E. (1955). “Relations entre la composition des familles et le taux demasculinité.” Journal de la société statistique de Paris 96: 49–64.

Mitter, N. S., and S. Anand (1975). “Trend Towards Unisexuality In Human Sibships : A Study In Two Generations.” Indian Anthropologist 5(2): 12–20.

Nagamani, M., P. G. McDonough, J. O. Ellegood and V. B. Mahesh (1979). “Maternal and amniotic fluid steroids throughout human pregnancy.” Am J Obstet Gynecol 134(6): 674–680.

Renkonen, K. O. (1956). “Is the sex ratio between boys and girls correlated to the sex of precedent children?” Ann Med Exp Biol Fenn 34(4): 447–451.

Renkonen, K. O., O. Makela and R. Lehtovaara (1961). “Factors affecting the human sex ratio.” Ann Med Exp Biol Fenn 39: 173–184.

Renkonen, K. O., O. Makela and R. Lehtovaara (1962). “Factors affecting the human sex ratio.” Nature 194: 308–309.

Rodgers, J. L., and D. Doughty (2001). “Does Having Boys or Girls Run in the Family?” CHANCE 14(4): 8–13.

Thomas Gualtieri, C., R. E. Hicks and J. P. Mayo (1984). “Influence of sex of antecedent siblings on the human sex ratio.” Life Sciences 34(19): 1791–1794.

Turpin, R. and M. P. Schutzenberger (1949). “Sur la distribution de sexes chez l’homme.” La Semaine des Hospitaux de Paris, La Semaine Medicale(25 annee, n. 60): 2544–2545.

Wikipedia. (2015). “Polygamy#Latter_Day_Saint_Movement.”

